# Current Situation of Breeding Bird Species in the Gediz Delta, Turkey

**DOI:** 10.1101/2022.06.11.494948

**Authors:** Şafak Arslan, Ahmet Kaya, Adem Akyol, Mehmet Kaya

## Abstract

Gediz Delta, one of the most important wetland ecosystems in the Mediterranean Basin, also is one of the most important areas for birds in Turkey. Breeding bird atlas studies were carried out in 2002, and 2006 and extensive breeding species were revealed. In 2021, we conducted an atlas of breeding birds using the same method as previous studies and observed the change over the last 15 years. Bird species seen and heard between May 3 and June 15 in 294 1x1 square kilometres were recorded and classified with internationally accepted breeding codes. Breeding codes were given to 113 species belonging to 48 families throughout the study. Of these, 32 species are classified as possible breeding, 34 species as probable breeding and 47 species as confirmed breeding. Threat factors of species and habitats are grouped under five main topics. According to previous studies, we observed that the number of breeding bird species increased. It is thought that the number of breeding bird species will increase with the elimination of threats and carrying out the necessary restoration works.

## 1. Introduction

Breeding bird atlas researches are important studies in which the current and abundance of bird species in a specific region are obtained, and these findings are poured onto the map. The distribution of bird species, their populations, their reactions to environmental changes and the factors threatening the species can be determined by creating appropriate grids in Atlas studies and taking samples in a specific period. Atlas studies are one of the most widely used methods for assessing the biodiversity of a region (Boyla et al., 2019). Turkey has different habitats thanks to its geographical location (Eken et al., 2006). In this way, bird species that need different requests are also seen in Turkey. There are 495 recorded bird species in Turkey so far (eBird, 2022). On the other hand, Boyla et al. (2019) took the regular breeding records of 313 species and the single breeding records of 3 species in Turkey, and a total of 316 bird species were given a breeding code. Gediz Delta one of the 305 Key Biodiversity Areas (KBA) in Turkey (Eken et al., 2006), meets the criteria of “Important Bird Area” for 28 bird species (Kılıç and Eken, 2004). Earliest information on the birds in the Gediz Delta date back to the middle of the 19th century (Gonzenbach, 1859; Krüper, 1869, 1875). Sıkı (1985) identified 182 bird species in his study, which was limited to the protected area of the Gediz Delta. The delta has been recorded as the breeding ground of a significant part of the breeding populations of Mediterranean gull, Caspian tern, sandwich tern and common tern species in the entire Mediterranean and where the first breeding colony of the sandwich tern in Turkey was discovered (Eken, 1997). In 2002, it was reported that 211 bird species and 59 of these species breed in the delta (Sıkı, 2002). In the first comprehensive breeding bird atlas on the Gediz Delta, 92 species were given a breeding code (Onmus et al., 2009). A similar study was repeated in 2006 and breeding codes were given to 104 bird species (Onmuş and Sıkı, 2010). The last breeding bird atlas study in the delta was carried out in 2014 (Ömer Döndüren, unpublished report). There was extensive studies on some bird groups in the Gediz Delta. Çiftçi (2006) worked on white stork, Gül (2014) on dalmatian pelicans, Döndüren (2015) on white stork and Kaya (2017) on tern species.

In our study, which was carried out comprehensively in two teams during the breeding period of birds, i) update the list of breeding bird species in the Gediz Delta and its current status, ii) compare previous studies and evaluate the land use of some species, iii) detect the threats.

## 2. Materyal Methods

### 2.1. Study Area

Gediz Delta (38° 30’N, 26° 55’E) is situated in the Western Aegean, covering approximately 40.000 ha. This delta is the largest on the Western Anatolian coasts, and it is known as the fourth largest delta of Turkey (Eken et al., 2006; Kaya, 2017). Gediz Delta includes salty, fresh and brackish water ecosystems. Most of the delta-sea border consists of sand bands covered with glasswort (*Salicornia* sp.) and mussel shells. Behind the sand bands take place lagoons or wide salt water coastal meadows. *Arthrocnema-Halocnemetum strobilaceum* formations in the coastal part of salt meadows, tamarisk (*Tamarix sp*) and *Limonium spp.* communities are included. At points where freshwater inflows into the salt area are high, there are small reeds and temporary wet meadows covered with rushes (*Juncus spp.*). The hills are usually covered with garrigue and scrub. Apart from this, there are large agricultural areas, plantation areas and gardens in the delta. One of the most important agricultural areas on the Aegean coast, the delta, especially the part known as Menemen Plain, has extremely fertile agricultural lands.

Our study did not include some of the large agricultural areas because they were similar. Thus, our total working area has been determined as 29,400 hectares.

### 2.2. Data Collection

29,400 ha of the study area is divided into 1×1 UTM (Universal Transverse Mercator) squares. The study was performed on 294 UTM frames (Figure 1). In each square, three points were taken to represent different habitats and to be 300 meters away from each other. Observations were made for 10 minutes at the determined counting points. All bird species observed and heard during this period were recorded. Breeding codes are given for birds in their breeding habitats (Onmuş et al., 2009; (Onmuş ve Sıkı, 2010). Species not found in their breeding habitats have not been recorded.

**Figure 1.**
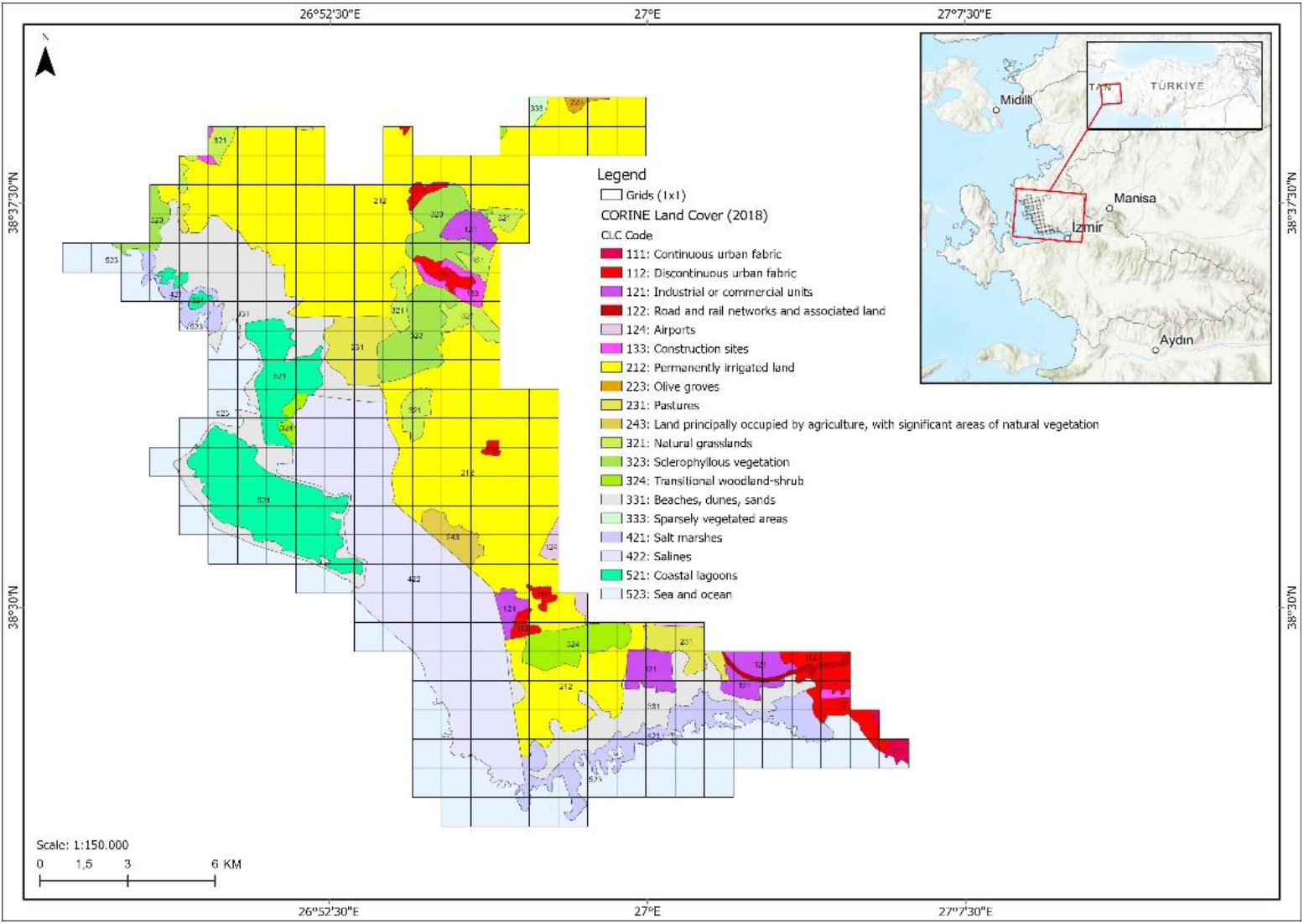
Study area divided into 1×1 km^2^ and habitat types

**Figure 2.**
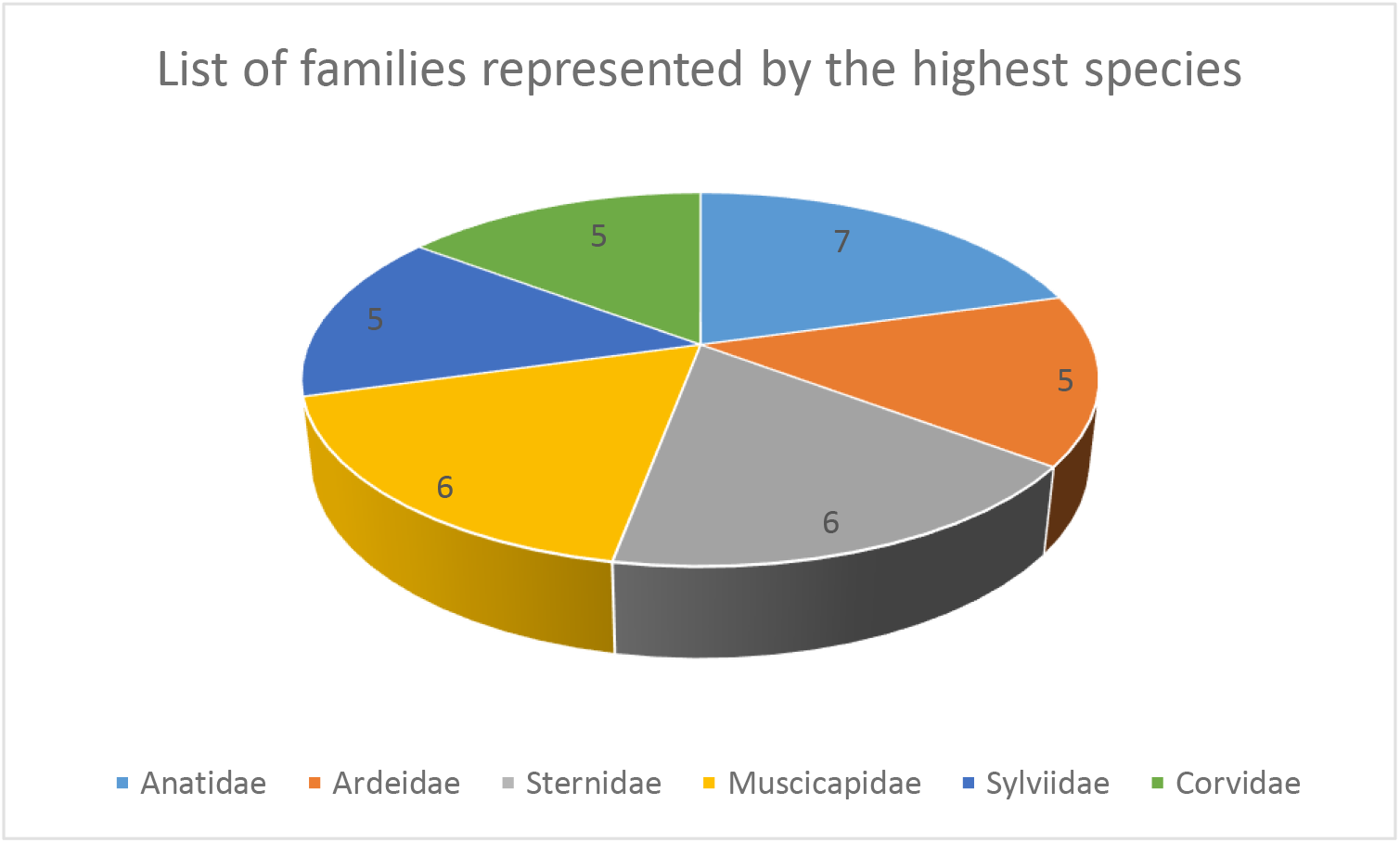
Families that included the highest species

Our study was carried out in two simultaneous teams between May 3 and June 15, which includes the breeding activity period of birds. During the observations, internationally accepted breeding codes established by the European Bird Census Committee (EBCC) were used. These codes are 16 in total and are divided into three main classes A (Possible breeding), B (Probable breeding) and C (Confirmed breeding), and these codes are given for each bird species according to the breeding behaviours of the birds (Keller et al., 2020).

Threat factors were also marked in forms at each point during the field studies. In this context, existing threats in the area were also collected. All the data collected during the fieldwork were digitized in excel format on the same day. In this way, the loss of data is also prevented.

Separate field studies were carried out for bird species as colonial breeding birds. The number of breeding pairs was determined by telescope and binoculars from the area where the breeding colonies could be seen and counted. For the colonial species that breed on the islets and on the coasts, censuses were carried out by the sea with the help of boats.

## 3. Results

During the breeding bird atlas study of the Gediz Delta, 143 bird species were observed, and 113 species (%79.02) were given a breeding code. Of these bird species were classified 32 possible breeding (%29.09), 34 probable breeding (30.9) and 47 confirmed breeding (%41.59) codes. 48 families represent these 113 species.

> *Botaurus stellaris; Ixobrychus minutus; Nycticorax nycticorax; Spatula querquedula; Falco tinnunculus; Falco subbuteo; Rallus aquaticus; Gelochelidon nilotica; Streptopelia turtur; Caprimulgus europaeus; Apus pallidus; Coracias garrulous; Upupa epops; Alauda arvensis; Motacilla alba; Luscinia megarhynchos; Oenanthe hispanica; Turdus philomelos; Acrocephalus palustris; Sylvia crassirostris; Sylvia curruca; Sylvia melanocephala; Sylvia ruppeli; Muscicapa striata; Poecile lugubris; Sitta neumayer; Oriolus oriolus; Lanius minor; Lanius nubicus; Garrulus glandarius; Chloris chloris; Emberiza schoeniclus* are classified as possible breeding.
>
> *Tachybaptus ruficollis; Podiceps cristatus; Mycrocarbo pygmeus; Ardeola ralloides; Ardea purpurea; Tadorna tadorna; Aythya farina; Aythya nyroca; Circus aeruginosus; Alectoris chukar; Recurvirostra avosetta; Tringa tetanus; Chlidonas hybrid; Columba livia; Clamator glandarius; Apus apus; Melanocorypha calandra; Cecropis daurica; Delichon urbicum; Cercotrichas galactotes; Turdus merula; Cettia cetti; Cysticola juncidis; Acrocephalus arundinaceus; Iduna pallida; Sylvia comminis; Lanius collurio; Corvus cornix; Corvus corax; Passer montanus; Fringilla coelebs; Carduelis carduelis; Emberiza caesia; Emberiza melanocephala* are classified as probable breeding.
>
> *Pelecanus crispus; Ciconia ciconia; Phoenicopterus roseus; Cygnus olor; Tadorna ferruginea; Anas platyrhynchos; Falco naumanni; Falco peregrinus; Gallinula chloropus; Fulica atra; Haematopus ostralegus; Himantopus Himantopus; Burhinus oedicnemus; Glareola pratincola; Charadrius alexandrines; Vanellus spinosus; Ichthyaetus melanocephalus; Larus genei; Larus michahellis; Hydroprogne caspia; Thalasseus sandvicensis; Sterna hirundo; Sternula albifrons; Streptopelia decaocto; Tyto alba; Athene noctua; Merops apiaster; Dendrocopus syriacus; Calandrella brachydactyla; Galerida cristata; Hirundo rustica; Anthus campestris; Motacilla flava; Saxicola torquatus; Oenanthe isabellina; Oenanthe oenanthe; Acrocephalus scirpaceus; Panurus biarmicus; Parus major; Remiz pendulinus; Lanius senator; Pica pica; Corvus monedula; Sturnus vulgaris; Passer hispaniolensis; Passer domesticus; Emberiza calandra* are classified as confirmed breeding.

When the habitat types of 1×1 km^2^ 294 squares were examined in the study area, permanently irrigated land, sea and ocean and salines were determined as the three dominant habitat types. Olive groves, sparsely vegetated areas and continuous urban fabric habitats were the habitats with the lowest density (Table 1). Permanently irrigated land habitat includes agricultural lands and salt production areas within the delta (Figure 1).

**Table 1.** Habitat type proportions of the study area

| CLC Code | CLC Name | Area (HA) | Area (%) |
| --- | --- | --- | --- |
| 223 | Olive groves | 36,19 | 0,12% |
| 333 | Sparsely vegetated areas | 47,12 | 0,16% |
| 111 | Continuous urban fabric | 62,99 | 0,21% |
| 122 | Road and rail networks and associated land | 84,09 | 0,29% |
| 124 | Airports | 107,92 | 0,37% |
| 133 | Construction sites | 167,81 | 0,57% |
| 243 | Land principally occupied by agriculture, with significant areas of natural vegetation | 199,68 | 0,68% |
| 324 | Transitional woodland/shrub | 466,94 | 1,59% |
| 321 | Natural grassland | 662,52 | 2,25% |
| 112 | Discontinuous urban fabric | 679,27 | 2,31% |
| 231 | Pastures | 714,48 | 2,43% |
| 121 | Industrial or commercial units | 740,68 | 2,52% |
| 323 | Sclerophyllous vegetation | 1.051,11 | 3,58% |
| 421 | Salt marshes | 1.103,33 | 3,75% |
| 521 | Coastal lagoons | 1.978,20 | 6,73% |
| 331 | Beaches, dunes, sands | 2.936,09 | 9,99% |
| 422 | Salines | 3.964,48 | 13,48% |
| 523 | Sea and ocean | 4.704,26 | 16,00% |
| 212 | Permanently irrigated land | 9.692,42 | 32,97% |
| **Total** |  | **29399,58** | **100,00%** |

### 3.1. Threats

The factors that threaten the birds in the Gediz Delta are grouped under five topics.

- It has been observed that the polluted waters of the industries and domestic wastes flow into the river feeding the delta. On the coasts, wastes mainly of glass and plastic, brought to the side by the waves or left by people, were also observed.
- Normally hunting is banned in the Gediz Delta, but hunting cartridges and duck blinds were seen in the study area. Poaching poses great pressure on duck species in particular.
- Although the delta has national and international protection boundaries, urbanization and mining activities pose pressure on the delta. The pressure of urbanization has decreased as of the 21st century, and mining activities are desired to be carried out, especially under the name of renewable energy. However, any mining activity is not happen in the current studies.
- Unplanned vehicle entry in the delta poses a significant threat. Especially the stable roads passing over birds’ breeding areas destroy nesting areas. It has been observed that there is more than one path reaching a point close to each other.
- For the irrigated agricultural activities carried out along the Gediz river that forms and feeds the delta, water is drawn with pumps from the Gediz river. For this reason, it has been determined that the freshwater areas in the delta are dry in the summer period.

When the threats in the Gediz Delta are examined, it is seen that the threats can be solved with simple planning and increasing the controls.

## 4. Discussion

In the 2021 Breeding Bird Atlas study, 113 bird species were given a breeding code, while 92 species were assigned a breeding code in 2002 (Onmuş et al., 2009) and in 2006, 104 species were assigned the breeding code (Onmuş and Sıkı, 2010). In 2002, out of 92 bird species, 47 species were given confirmed breeding (C) status, 22 species were assessed to be probably breeding (B) and 23 species were given possible breeding (A) codes. In 2006, out of 104 bird species, 61 species were assigned the code C, 25 species were assigned the code B and 17 species were assigned the code A. In 2021, 47 species of 113 species were given the code C, 35 species were given the code B and 31 species were given the code A. A comparison of the given breeding codes by year is given in table 2.

**Table 2.** Distribution of breeding codes by year

| Year<br>Code | 2002 | 2006 | 2021 |
| --- | --- | --- | --- |
| A | 23 | 17 | 31 |
| B | 22 | 25 | 35 |
| C | 47 | 61 | 47 |

The little egret (*Egretta garzetta*), Eurasian spoonbill (*Platalea leucorodia*), little-ringed plover (*Charadrius dubius*), black tern (*Chlidonias niger*), common cuckoo (*Cuculus canorus*), and the white-throated robin (*Irania guttularis*) were given breeding codes only in 2002, they could not be given in 2021. While it is known that the little egret established an incubation colony in the delta in the 1980s, it was not observed to be incubating in later years (Eken, 1997).

The Ruddy turnstone (*Arenaria interpres*), Eurasian eagle-Owl (*Bubo bubo*), alpine swift (*Apus melba*), Eurasian blackcap (*Sylvia atricapilla*), common chiffchaff (*Phylloscopus collybita*), Eurasian blue tit (*Cyanistes caeruleus*), and cirl bunting (*Emberiza cirlus*) were given a breeding codes in 2006, but could not be given to them in 2021.

The Montagu’s harrier (*Circus pygargus*) and sand martin (*Riparia riparia*) were given breeding codes in both 2002 and 2006, but could not be given it in 2021.

The great crested grebe (*Podiceps cristatus*), pygmy cormorant (*Microcarbo pygmeus*), Eurasian bittern (*Botaurus stellaris*), black-crowned night heron (*Nycticorax nycticorax*), mute swan (*Cygnus olor*), common pochard (*Ayhtya ferina*), ferruginous duck (*Aythya nyroca*), Eurasian hobby (*Falco subbuteo*), song thrush (*Turdus philomelos*), Eastern orphean warbler (*Sylvia crassirostris*), Rüppell’s warbler (*Sylvia ruppeli*), Eurasian golden oriole (*Oriolus oriolus*), Northen raven (*Corvus corax*) and the Cretzschmar’s bunting (*Emberiza caesia*) were given breeding codes in 2021, but was not given them in 2002 or 2006.

Of the 113 bird species given a breeding code in the atlas study, dalmatian pelican (*Pelecanus crispus*), ferruginous duck (*Aythya nyroca*) and Eurasian oystercatcher (*Haematopus ostralegus*) while they are classified in the near threatened (NT) category, common pochard (*Aythya ferina*) and turtle dove (*Streptopelia turtur*) is classified in the vulnerable (VU) endangered category based on the red list categories of The International Union for Conservation of Nature (IUCN). The other 108 species are in the least concern (LC) category.

Our study conducted in 2021 shows that there is an increase in the number of breeding bird species in the Gediz Delta. However, the main threats to the species and the delta continue. Although we have classified the threats into five topics during our study, it is seen that the issues that need to be resolved urgently and quickly are drying up due to illegal hunting, wrong water and irrigated agricultural policies. If these threats are eliminated, it is thought that the continuity of suitable habitats for the species will be ensured in the Gediz Delta, and an increase in the number of species will occur over time.

## Acknowledgements

This study was carried out with the permission of the General Directorate of Nature Conservation and National Parks and with the contribution of Izmir Metropolitan Municipality.

## References

Boyla, K.A., Sinav, L. ve Dizdaroğlu D.E. 2019. Türkiye Üreyen Kuş Atlası. WWF-Türkiye, Doğal Hayati Koruma Vakfi. Istanbul.

Çiftçi, A. (2006). İzmir’de Leylek *Ciconia ciconia* (L., 1758) popülasyonunun Tespiti üzerine Araştirmalar. Yüksek Lisans Tezi. Biyoloji Anabilim Dalı, Fen Bilimleri Enstitüsü, Ege Üniversitesi, izmir. 60 syf. (in Turkish Thesis)

Döndüren, Ö. (2015). Gediz Nehri Havzası’nda Leylek *(Ciconia ciconia* L., 1758) Popülasyonunun Tespiti, Popülasyon Değişimlerinin ve Popülasyonu Etkileyen Etmenlerin Belirlenmesi ile Türün Göç Dinamikleri Üzerine Araştirmalar. Doktora Tezi, Ege Üniversitesi, İzmir. (in Turkish Thesis)

eBird, Birds of Turkey List. https://ebird.org/region/TR. 12.05.2022.

Eken G, Bozdoğan M, Isfendiyaroğlu S, Kiliç DT, Lise Y (2006). Türkiye’nin Önemli Doğa Alanlari. Doğa Derneği, Ankara (in Turkish).

Eken, G., 1997. The Breeding Population of some Species of Waterbirds at Gediz Delta, Western Turkey. Zoology in the Middle East, Vol. 14:53–68. Max Kasparek Verlag, Heidelberg.

Gonzenbach, G. V., (1859): Excursionen an die Brutplätze von Sterna, Larus und Glareola im Golf von Smyrna im Frühling 1859. — Journal für Ornithologie 7: 308–316, 393–398, Leipzig.

Gül, O. (2014). Gediz Deltasi (izmir) ve Büyük Menderes Deltasi (Aydin)’nda Üreyen Tepeli Pelikan *(Pelecanus crispus* Bruch, 1832) Populasyon Büyüklüğünün, Değişiminin, Göçlerinin, Üreme ve Beslenme Biyolojilerinin Araştirilmasi. Doktora Tezi, Ege Üniversitesi, İzmir. (in Turkish Thesis)

Kaya, A. (2017). Homa Dalyani Yaban Hayati Koruma Sahasinda Kuluçkaya Yatan Kuş Türlerinden Yapay Üreme Platformlarindaki Kuluçkaya Yatan Kuş Türlerinin Tespiti ve Üreme Başarilarinin Araştirilmasi. Yüksek Lisans Tezi, Biyoloji Anabilim Dalı, Fen Bilimleri Enstitüsü, Ege Üniversitesi, İzmir. (in Turkish Thesis)

Keller, V., Herrando, S., Voríšek, P., Franch, M., Kipson, M., Milanesi, P., … & Foppen, R. P. B. (2020). European breeding bird atlas 2: Distribution, abundance and change.

Kiliç DT, Eken G. 2004. Türkiye’nin Önemli Kuş Alanları - 2004 güncellemesi (Important Bird Areas in Turkey - 2004 update). Ankara: Doga Dernegi.

Krüper, Th. (1869): Beitrag Zur Ornithologie Kleinasiens. - Journal Fllr Ornithologie 17: 21–45, Leipzig.

Krüper, Th. (1875): Beitrag Zur Ornithologie Kleinasiens. - Journal Fllr Ornithologie 23: 258–285, Leipzig.

Onmuş, O. ve Sıkı, M., 2010. State of the Breeding Birds in Gediz Delta: Distributions, Abundances, and Changes in Bird populations. Bird Census News 2010, 23/1-2: 59–69.

Onmuş, O., Durusoy, R. ve Eken, G., 2009. Distribution of breeding birds in the Gediz Delta, Western Turkey. Zoology in the Middle East 47: 39–48. Max kasparek Verlag, Heidelberg.

Siki, M. (1985). Çamalti Tuzlasi - Homa Dalyani Kuş Türleri ve Bazı Türlerin Biyolojileri Üzerine Araştirmalar. Ege Üniversitesi Fen Fakültesi Tabiat Tarihi Müzesi, İzmir. (in Turkish Thesis)

Siki, M. (2002). Gediz Deltasi (Izmir Kuş Cenneti) Kuşları. Ekoloji Çevre Dergisi. Cilt:11, Sayı:44, 11–16 (in Turkish).

